# Exploration of novel biomarkers for neurodegenerative diseases using proteomic analysis and ligand-binding assays

**DOI:** 10.1101/2024.11.12.622994

**Authors:** Annalena Kenzelmann, Christina Boch, Ronny Schmidt, Mario Richter, Michael Schulz

## Abstract

Neurodegenerative diseases are a major cause of morbidity and mortality world-wide, and their public health burden continues to increase. There is an urgent need to develop reliable and sensitive biomarkers to aid the timely diagnosis, disease progression monitoring, and therapeutic development for neurodegenerative disorders. Proteomic screening strategies, including antibody microarrays, are a powerful tool for biomarker discovery, but their findings should be confirmed using quantitative assays. The current study explored the feasibility of combining an exploratory proteomic strategy and confirmatory ligand-binding assays to screen for and validate biomarker candidates for neurodegenerative disorders. It analyzed cerebrospinal fluid (CSF) and plasma samples from patients with Alzheimer’s disease, Parkinson’s disease, and multiple sclerosis and healthy controls. The screening antibody microarray identified differentially expressed proteins between patients with neurodegenerative diseases and healthy controls. Quantitative ligand-binding assays confirmed that cluster of differentiation 14 (CD14) levels were elevated in CSF of patients with Alzheimer’s disease, whereas osteopontin levels were increased in CSF of patients with Parkinson’s disease. The current study demonstrated the utility of combining an exploratory proteomic approach and quantitative ligandbinding assays to identify biomarker candidates for neurodegenerative disorders. To further validate and expand these findings, large-scale analyses using well characterized samples should be conducted.

## Introduction

Neurodegenerative diseases are an important cause for morbidity, premature mortality, and public health burden globally, especially among older adults [1,2]. Since life expectancy is increasing, and age is the most important risk factor for neurodegenerative diseases, their incidence continues to rise [2,3].

Neurodegenerative diseases are characterized by irreversible and progressive neuronal dysfunction and loss, leading to a progressive clinical course and cognitive and motor dysfunction [3,4]. Several of the neurodegenerative diseases with highest prevalence and public health impact include Alzheimer’s disease, Parkinson’s disease, multiple sclerosis, and amyotrophic lateral sclerosis [4,5].

The pathophysiology of neurodegeneration is complex and heterogenous and involves multiple pathways. Pathological aggregates and oligomers of neurodegenerative disease-associated proteins have been detected in patients with neurodegenerative disorders, including amyloid-β, tau, α-synuclein, and transactive response DNA-binding protein 43 (TDP-43) [6,7]. In addition, proteotoxic and oxidative stress, neuroinflammation, abnormalities in the autophagosomal and ubiquitin-proteosome system, and apoptosis have been implicated in the pathophysiology of neurodegenerative diseases [3,4].

As neurodegenerative diseases are often characterized by an asymptomatic prodromal phase, biomarker development may facilitate their early and precise diagnosis, which may also have implications for therapy [8,9]. In addition, biomarkers may aid the identification of therapeutic targets and monitoring of disease progression and therapeutic responses [9–11].

Molecules related to pathophysiological mechanisms implicated in neurodegenerative diseases, like neuronal loss associated with protein deposits in the brain, can be promising biomarker candidates detected with imaging modalities or in cerebrospinal fluid (CSF) or blood [3,7,10]. In CSF, neurofilament light chain (NfL) and total tau (t-tau) have emerged as markers of axonal degeneration observed in neurodegenerative diseases [12]. Moreover, phosphorylated tau (p-tau), amyloid-β, alpha-synuclein, and TDP-43 have shown promise as CSF biomarkers reflecting a particular type of pathological protein accumulation [12]. In blood, the concentrations of amyloid-β and p-tau have also emerged as markers of amyloid and tau pathologies characteristic of Alzheimer’s disease [13]. In addition, blood levels of NfL, ubiquitinC-terminal-hydrolase-L1, and β-synuclein have been identified as markers of neuronal degeneration, whereas glial fibrillary acidic protein has been proposed as a marker of glial degeneration [13]. Despite these research advances, there is still an urgent need to develop sensitive and reliable biomarkers for neurodegenerative disorders [3].

The identification of CSF biomarkers for neurodegenerative disorders is of particular interest, because CSF is the biofluid located closest to brain structures and may thus be most likely to reflect pathological brain changes [14]. However, since performing a lumbar puncture to withdraw CSF is associated with patient discomfort, longitudinal CSF sampling and disease progression monitoring are challenging. In contrast, blood is a more easily accessible biofluid for longitudinal sample collection. An ideal biomarker for neurodegenerative disorders would show the same pattern in CSF and blood and would reflect pathophysiological changes in the brain.

Omics-based technologies are a powerful tool for biomarker identification as they enable the simultaneous screening of numerous molecules for their differential expression in individuals affected by different medical conditions, including neurodegenerative diseases. While genomic or transcriptomic techniques have repeatedly been employed for biomarker screening, technological advances have also enabled proteomic screening strategies that analyze a wide range of functional molecular products with various biological functions [15]. Antibody microarrays are proteomic screening tools that have gained popularity due to their sensitivity, reproducibility, high-throughput capabilities, and convenience of use [15–17]. However, the findings of protein microarrays should be further validated using quantitative assays, such as ligand-binding assays. Antibody microarrays have been successfully implemented for the identification of biomarker candidates in several disease areas, including gastric cancer and endometriosis [15,18,19]. Nevertheless, there is scarcity of studies combining antibody microarrays with ligand-binding immunoassays for the identification of biomarkers for neurodegenerative disorders.

The current study aimed to explore the feasibility of combining an exploratory antibody microarray and validatory ligand-binding assays to identify biomarker candidates in CSF and plasma samples of patients with neurodegenerative disorders (Alzheimer’s disease, Parkinson’s disease, multiple sclerosis, and amyotrophic lateral sclerosis) compared to healthy controls.

## Materials and Methods

### Sample selection

Plasma and CSF samples from healthy controls and from patients with Alzheimer’s disease, Parkinson’s disease, multiple sclerosis, and amyotrophic lateral sclerosis were purchased from BioIVT/PrecisionMed, Biotrend, Central BioHub, and Discovery Life Sciences.

### Proteomic screening with the ScioDiscover platform

Proteomic screening was performed using scioDiscover, a chip-based microarray system that profiles samples for changes in 1,351 proteins (Sciomics GmbH, Neckargemünd, Baden-Württemberg, Germany). The scioDiscover platform includes immobilized antibodies against a variety of targets, such as chemokines, cytokines, growth factors, immune cell markers, immune checkpoint molecules, inflammatory mediators, and vascular/angiogenic targets, and can be used for the analysis of samples derived from CSF, plasma, cells, tissues, supernatants, and organoids. The workflow of the scioDiscover screening platform is illustrated in Figure 1.

**Figure 1.**
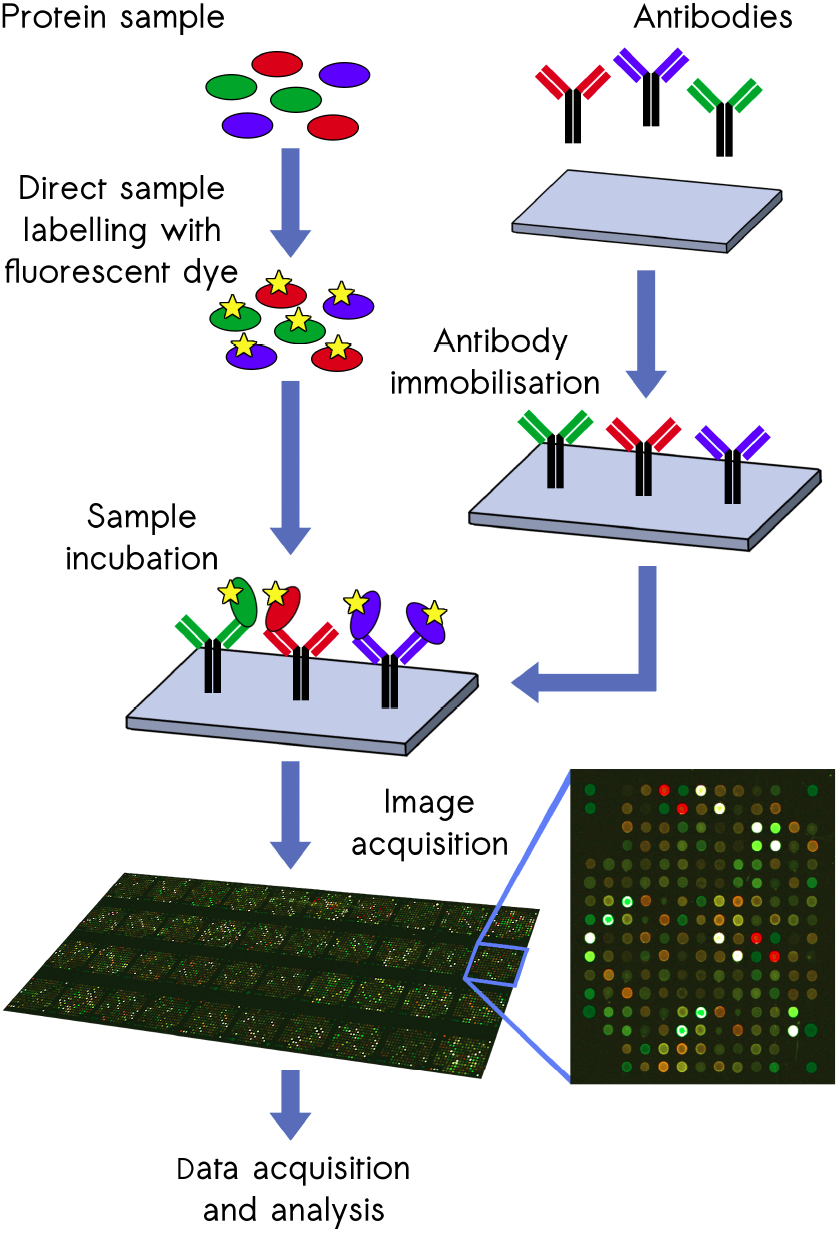
Antibody microarray experiment pipeline.

First, proteins were extracted, and the bulk protein concentration was determined by BCA assay. A reference sample was established by pooling an identical volume of each sample.

CSF and plasma samples were labeled at adjusted protein concentrations of 4 mg/ml for plasma and 0.6 mg/ml for CSF for 2 hours with scioDye 1. The reference sample was labeled with scioDye 2. After 2 hours, the reaction was terminated, and the buffer was exchanged to phosphate-buffered saline (PBS). All labelled protein samples were stored at -20 °C until further use.

Samples were analyzed using a dual-color approach and a reference-based design on scioDiscover antibody microarrays (Sciomics) targeting 1,351 different proteins with 1,821 antibodies. Each antibody was represented on the array in four replicates. The arrays were blocked with scioBlock (Sciomics) on a Hybstation 4800 (Tecan, Austria), and afterwards the samples were incubated competitively with the reference sample using a dual-color approach. After incubation for 3 hours, the slides were thoroughly washed with 1 ×PBSTT, rinsed with 0.1 ×PBS as well as with water, and dried with nitrogen.

Slide scanning was conducted with a PowerScanner (Tecan, Austria) with constant instrument laser power and PMT settings. Spot segmentation was performed with GenePix Pro 6.0 (Molecular Devices, Union City, CA, USA).

### Statistical analysis of the proteomic screening data

Acquired raw data were analyzed using the linear models for microarray data (*limma*) package of R-Bioconductor after uploading the median signal intensities. For normalization, a specialized invariant LOWESS method was applied. For sample analysis, a one-factorial linear model was fitted with *limma*, resulting in a two-sided t-test or F-test based on moderated statistics. All presented p values were adjusted for multiple testing by controlling the false discovery rate according to Benjamini and Hochberg. Proteins were defined as differential for |logFC| > 0.5 and an adjusted p value < 0.05.

Differences in protein abundance between samples or sample groups are presented as logfold changes (logFC) calculated for basis 2. In a study comparing samples versus controls, a logFC = 1 means that the sample group had on average a 2^1^ = 2-fold higher signal than the control group. logFC = −1 stands for 2^−1^ = 1/2 of the signal in the sample as compared to the control group.

### Automated fluorescence immunoassays

Automated fluorescence Simple Plex™ immunoassays for single-analyte measurements were performed using the Ella™ Automated Immunoassay System (Bio-Techne, ProteinSimple, Minneapolis, MN, USA), which enables rapid, sensitive, and highly reproducible detection of bioanalytes in various biological matrices.

The Simple Plex™ Human CD14 (SPCKB-PS-000386) and Simple Plex™ Human OPN 2^nd^ gen (SPCKB-PS-004314) assays were used in the current study (Bio-Techne, ProteinSimple, Minneapolis, MN, USA). Diluted samples and wash buffer were added on a microfluidic assay cartridge, which included a preloaded capture antibody, detection antibody, and fluorescent detector. After the samples and washing buffer were loaded in sample triplicates, the cartridge was transferred into the Ella analyzer, which performed the automated pipetting steps and read out the signal. A 631 nm laser was employed to excite the detect fluor, and a CCD camera was used to read the fluorescence signal, which was proportional to the amount of analyte present in each sample [20].

### Standards, quality controls, and dilution linearity

Standards and quality controls were run with each cartridge of the quantitative ligandbinding assays. The acceptance criteria for coefficient of variation and bias were 20% (lower limit of quantification; LLOQ 25%) for standards and 30% for quality controls.

As a quality criterion, the assays were tested for dilution linearity. Individual samples from healthy controls were serially diluted and analyzed with the respective assay. The actual concentration of the analyte in the samples was then back-calculated, and this determined concentration had to be in a range between -/+ 30% of the mean value of the actual concentration for each dilution.

### Statistical analysis of the fluorescence immunoassay data

Concentrations were back-calculated and dilution-corrected. Statistical analyses were performed on data that were logarithmically transformed according to the equation Y = log10 Y.

If the different groups within a data set yielded a skewness in the range of -1 to 1, a normal distribution was assumed, and a parametric test was selected. In the case of parametric tests, for data sets without significantly different standard deviations (SDs), ordinary one-way analysis of variance (ANOVA) with Dunnett’s multiple comparisons test was performed, and otherwise Welch’s ANOVA with Dunnett’s T3 multiple comparisons test was carried out. For data sets with more than one group with skewness outside the range, a nonparametric Kruskal-Wallis test with Dunn’s multiple comparisons test was chosen. The confidence level was set at 95% for all tests.

For the CD14 fluorescence immunoassay, plasma data were statistically analyzed via parametric Welch’s ANOVA with Dunnett’s T3 multiple comparisons test, whereas CSF data were statistically analyzed via the non-parametric Kruskal-Wallis test with Dunn’s multiple comparison test. For the osteopontin fluorescence immunoassay, plasma data were statistically analyzed via parametric Welch’s ANOVA with Dunnett’s T3 multiple comparisons test, whereas CSF data were statistically analyzed via the non-parametric Kruskal-Wallis test with Dunn’s multiple comparison test.

For samples, which resulted in low relative fluorescence unit (RFU) values below the LLOQ, a respective value was included in the statistical analysis with 1/2 LLOQ (only one sample corresponded to these criteria).

Statistical analyses were performed using GraphPad Prism Version 9.3.1 (GraphPad, San Diego, CA, USA).

## Results

### Characteristics of the study participants

Proteomic screening was conducted on a discovery cohort, which included 20 CSF samples and 32 plasma samples. The discovery cohort encompassed five CSF samples and eight plasma samples from each of the following groups: healthy controls, patients with Alzheimer’s disease, patients with Parkinson’s disease, and patients with multiple sclerosis.

Validation of the screening findings with quantitative ligand-binding assays was performed on an expanded validation cohort, which also included the samples from the discovery cohort. The CSF samples in the validation cohort were obstained from 84 controls, 18 patients with Alzheimer’s disease, 35 patients with Parkinson’s disease, 10 patients with multiple sclerosis, and 14 patients with amyotrophic lateral sclerosis. The plasma samples in the validation cohort were obtained from 11 controls, 14 patients with Alzheimer’s disease, 13 patients with Parkinson’s disease, nine patients with multiple sclerosis, and five patients with amyotrohic lateral sclerosis. Further information on the demographic characteristics of the CSF and plasma samples from the validation cohort are presented in Supplementary Tables S1 and S2, respectively.

### Differentially expressed proteins in patients with neurodegenerative disorders detected using the scioDiscover proteomic screening platform

In the proteomic antibody microaray, for CSF samples, more than 90% of the antibodies exhibited a signal-to-noise ratio (SNR) above 2, whereas more than 85% of the antibodies exhibited a signal-to-background ratio (SBR) above 2. For plasma samples, more than 95% of the antibodies exhibited an SNR above 2 and SBR above 2. The coefficients of variation (CVs) were below 10% for the majority of features for both CSF and plasma samples. On average, the biological variation among CSF samples was 29.9%, whereas the biological variation among plasma samples was 55.5% (median, 43.0%).

The proteomic biomarker screening identified a number of differentially expressed proteins between patients with Alzheimer’s disease, Parkinson’s disease, or multiple sclerosis and healthy controls. For CSF samples, in comparison to healthy controls, the number of antibodies that recorded differential protein abundance was 115 in patients with Alzheimer’s disease, nine in patients with Parkinson’s disease, and one in patients with multiple sclerosis. A list of the most differentially expressed proteins (p < 1e-02) identified by the antibody microarray in CSF samples from patients with each neurodegenerative disease in comparison to controls is presented in Supplementary Table S3. The CSF findings of the proteomic screening for patients with Alzheimer’s disease and Parkinson’s disease in comparison to healthy controls are also presented in Figure 2.

**Figure 2.**
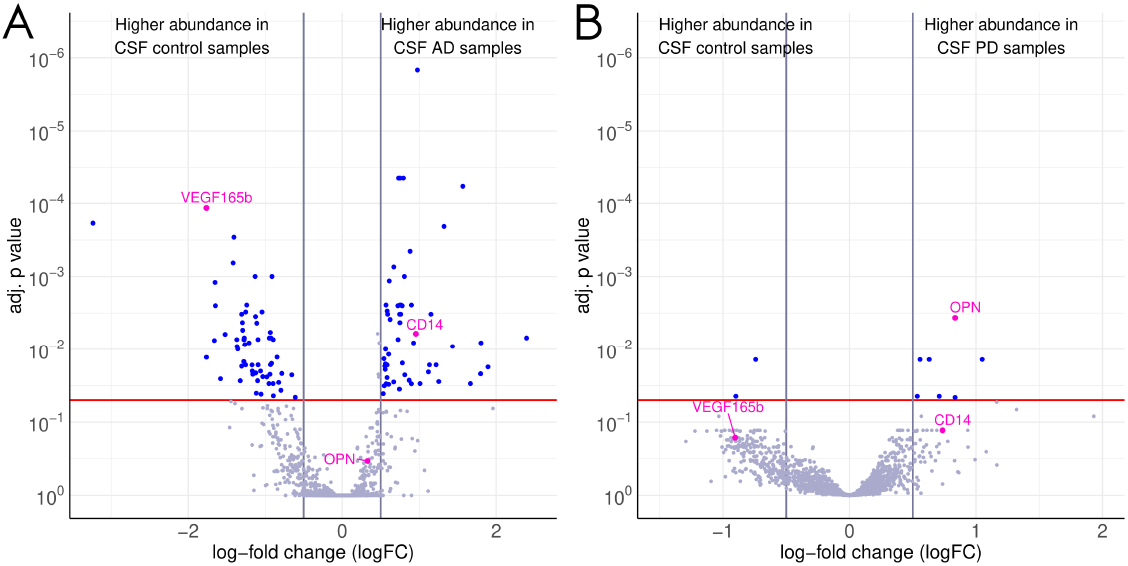
Differentially expressed proteins in CSF samples from patients with Alzheimer’s disease (**A**) and Parkinson’s disease (**B**) in comparison to healthy controls identified using the scioDiscover proteomic screening array. The red line indicates the significance cut-off of an adjusted p-value < 0.05.

For plasma samples, in comparison to healthy controls, the number of antibodies that detected differential protein abundance in comparison to controls was 21 in patients with Alzheimer’s disease, 30 in patients with Parkinson’s disease, and 12 in patients with multiple sclerosis. A list of the most differentially expressed proteins (p < e-02) identified by the antibody microarray in plasma samples from each neurodegenerative disease in comparison to controls is presented in Supplementary Table S4.

For the proteins, which were differentially expressed in patients with neurodegenerative diseases according to the proteomic screening assay, a literature search was conducted to examine their potential relevance, and the market availability of suitable immunofluorescence or chemiluminescence assays for their validation was examined. Our main goal was to evaluate the workflow combining an exploratory antibody microarray and a confirmatory ligandbinding assay rather than to investigate each potential biomarker. Finally, cluster of differentiation 14 (CD14), whose expression was increased in CSF from patients with Alzheimer’s disease (adjusted p = 5.9e-03) and plasma from patients with Parkinson’s disease (adjusted p = 3.4e-04), and osteopontin, whose expression was elevated in CSF from patients with Parkinson’s disease (adjusted p = 3.5e-03), were selected for validation with quantitative ligand-binding assays (Figures 2 and 3).

**Figure 3.**
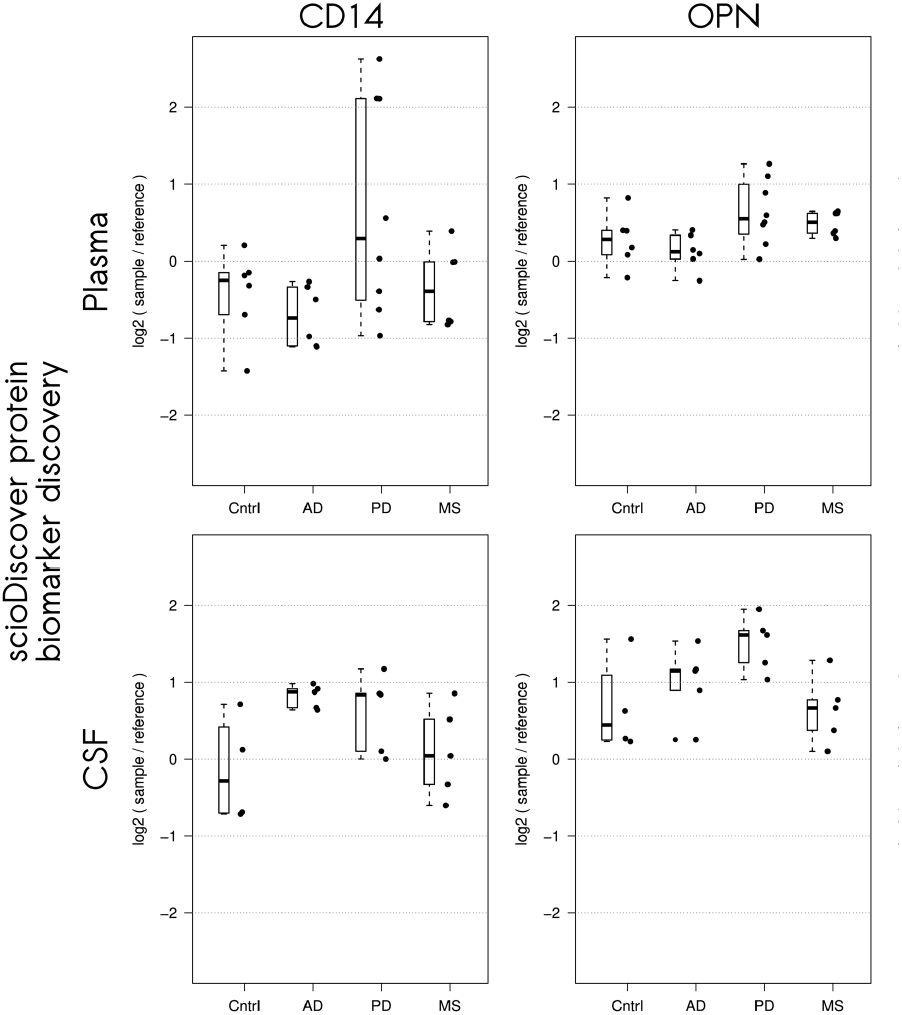
Abundance of cluster of differention (CD14) and osteopontn (OPN) identified using the scioDiscover platform in controls (Cntrl) and patients with Alzheimer’s disease (AD), Parkinson’s disease (PD), and multiple sclerosis (MS). Protein abundance was measured by the log-ratio of signals of individual samples and a reference sample.

### Validation of elevated CD14 levels in CSF of patients with Alzheimer’s disease using an automated fluorescence immunoassay

First, the dilution linearity of the Simple Plex™ Human CD14 assay was confirmed in CSF and plasma samples, using three different individual samples and a pooled sample containing CSF or plasma from at least three healthy individuals.

Next, CD14 protein expression was analyzed in plasma and CSF samples from patients with Alzheimer’s disease, Parkinson’s disease, multiple sclerosis, and amyotrophic lateral sclerosis and from healthy controls (dilution 1:2000 for plasma samples and 1:500 for CSF samples). Increased CD14 levels were detected in CSF samples from patients with Alzheimer’s disease in comparison to healthy controls (p = 0.0177). The CD14 levels in plasma samples from patients with Parkinson’s disease showed a tendency towards an increase in comparison to controls, which did not reach statistical significance (p = 0.0650) (Figure 4).

**Figure 4.**
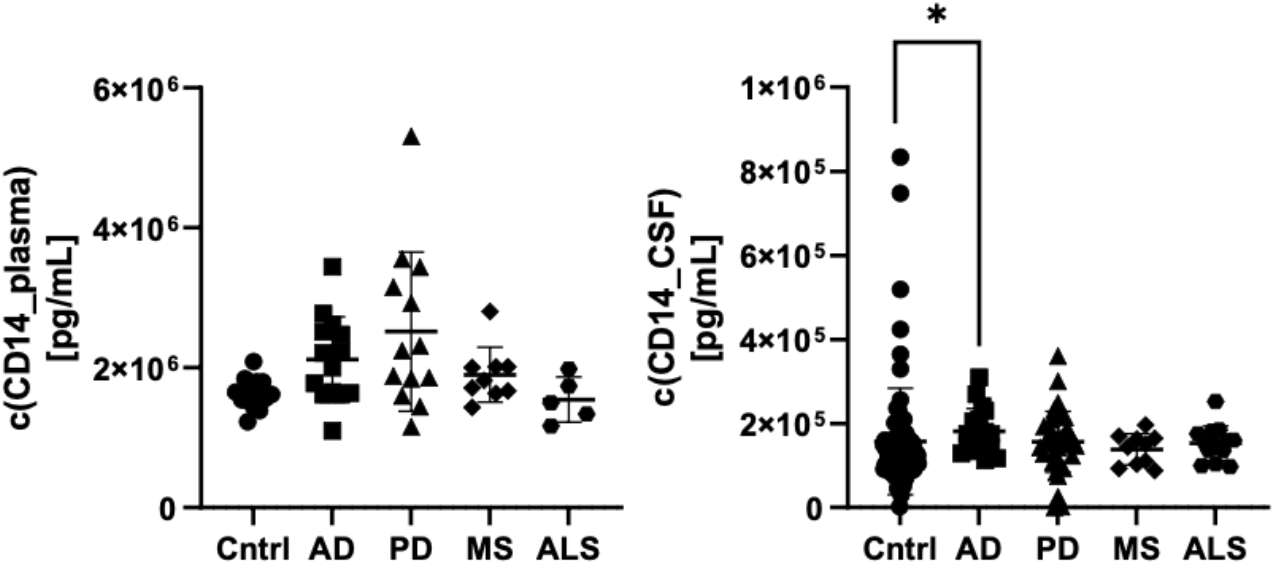
CD14 levels in plasma (left panel) and CSF (right panel) of patients with Alzheimer’s disease, Parkinson’s disease, multiple sclerosis, and amyotrophic lateral sclerosis and from healthy controls measured with the automated fluorescence-based Simple Plex™ Human CD14 assay. * p < 0.05. AD: Alzheimer’s disease; ALS: amyotrophic lateral sclerosis; CSF: cerebrospinal fluid; CD14: cluster of differentiation 14; Cntrl: controls; MS: multiple sclerosis; PD: Parkinson’s disease.

### Validation of elevated osteopontin levels in CSF of patients with Parkinson’s disease using an automated fluorescence immunoassay

The dilution linearity of the Simple Plex™ Human OPN assay in plasma and CSF was confirmed using serial dilutions of three individual samples and a pooled sample from healthy individuals for each matrix.

Osteopontin levels were measured in plasma and CSF samples from patients with Alzheimer’s disease, Parkinson’s disease, multiple sclerosis, and amyotrophic lateral sclerosis and from healthy controls using the automated fluorescence-based Simple Plex™ Human OPN assay. The fluorescence immunoassay validated the elevated expression of osteopontin in CSF from patients with Parkinson’s disease in comparison to CSF from healthy controls (p = 0.0346) (Figure 5). In addition, the fluorescence-based Simple Plex™ Human OPN assay observed increased osteopontin levels in plasma from patients with Parkinson’s disease (p = 0.0011) or multiple sclerosis (p = 0.0071) in comparison to healthy controls as well as decreased osteopontin concentration in CSF of patients with multiple sclerosis (p = 0.0219) in comparison to healthy controls (Figure 5).

**Figure 5.**
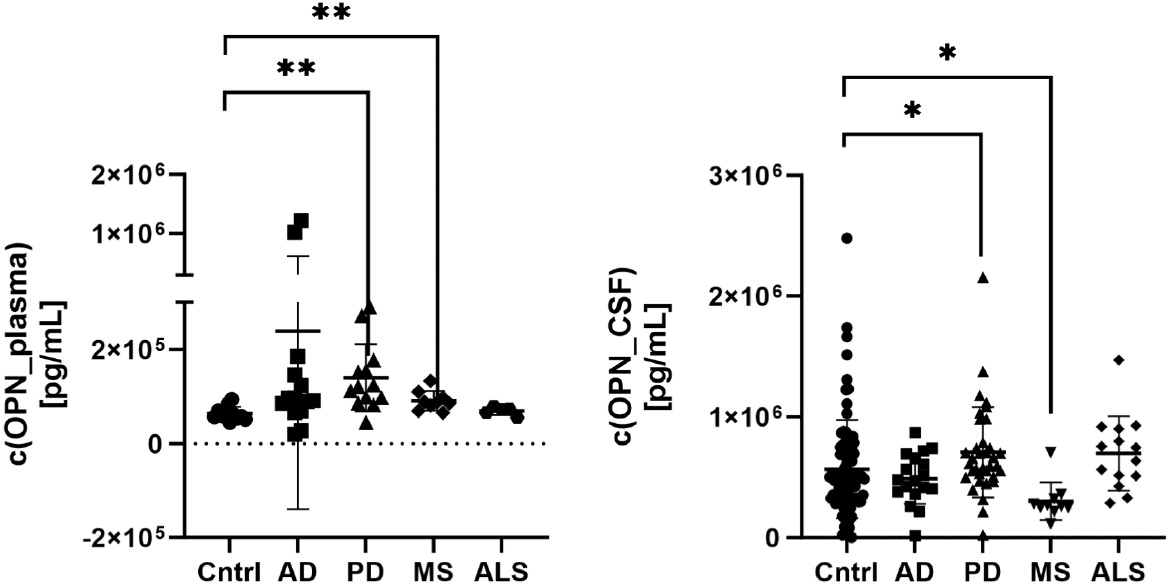
Osteopontin levels in plasma (left panel) and CSF (right panel) samples from patients with Alzheimer’s disease, Parkinson’s disease, multiple sclerosis, and amyotrophic lateral sclerosis and from healthy controls determined using the automated fluorescence-based Simple Plex™ Human OPN assay. * corresponds to p < 0.05 and ** to p < 0.01. AD: Alzheimer’s disease; ALS: amyotrophic lateral sclerosis; CSF: cerebrospinal fluid; Cntrl: controls; MS: multiple sclerosis; OPN: osteopontin; PD: Parkinson’s disease.

## Discussion

The current study successfully combined a proteomic antibody microarray and quantitative ligand-binding assays to screen for and validate differentially expressed proteins in patients with neurodegenerative disorders, including elevated CD14 in CSF from patients with Alzheimer’s disease and osteopontin in CSF from patients with Parkinson’s disease.

Proteomic assays have increasingly been used as screening tools for the identification of biomarker candidates, as they detect a wide range of functional molecular products with biological activity [15]. Antibody microarrays have been of particular interest, since they allow sensitive, reproducible, and high-throughput multiplexed analysis of proteins from a variety of sample types and in a convenient format [15,16]. However, the findings of antibody microarrays should be validated using quantitative ligand-binding assays. This combinational approach, even though used in other disease areas, has, to the best of our knowledge, not been established for the identification of biomarker candidates for neurodegenerative disorders. The current study demonstrated that a combination of the scioDiscover antibody microarray and ligand-bindings arrays can be used for identification of biomarker candidates in patients with neurodegenerative disorders.

Increased CD14 concentration was detected in CSF of patients with Alzheimer’s disease in comparison to controls by both the proteomic screening assay and quantitative ligand-binding assays.

CD14 is a pattern recognition receptor that has been implicated in inflammation, the innate immune response, metabolic disorders, and tumorigenesis and can hyperactivate microglia [21,22]. It exists in membrane-bound (expressed on various cell types, including monocytes, macrophages, microglia, and dendritic cells) and soluble (secreted by shedding and present in body fluids like CSF and serum) forms. CD14 sensitizes cells to lipopolysaccharide by transferring lipopolysaccharide molecules to its co-receptor Toll-like receptor 4 (TLR4). Interactions between CD14 and TLR4 mediate inflammation via production of proinflammatory cytokines [22].

CD14 has previously been implicated in Alzheimer’s disease. Amyloid-β plaques, which are a major feature of Alzheimer’s disease, induce the immune response via microglial activation and neuroinflammation. As a coreceptor of TLR2 and TLR4, CD14 binds to fibrillar amyloid-β [23]. Moreover, CD14 plays a role in amyloid-β-induced microglial activation and neurotoxicity and in the phagocytosis of amyloid-β [24,25]. Upregulated CD14 levels have been demonstrated in brain tissues of murine models of Alzheimer’s disease compared to control animals, and CD14-positive cells have been detected around amyloid-β plaques in post-mortem brains from patients with Alzheimer’s disease [26]. In addition, Busse et al. (2021) detected an increase in CD14+ monocytes in CSF of patients with mild and moderate Alzheimer’s disease [27]. These observations are in line with the finding of this study that CD14 was increased in CSF of patients with Alzheimer’s disease.

In the current study, the scioDiscover proteomic screening platform detected elevated CD14 levels in plasma samples from patients with Parkinson’s disease compared to controls. The quantitative CD14 fluorescence-based immunoassay identified a trend towards CD14 increase in plasma of patients with Parkinson’s disease, which did not reach statistical significance (p = 0.0650). Interestingly, this trend was only found in plasma and not in CSF. However, a previous study detected upregulated CD14 levels in post-mortem brain tissues of a murine model of Parkinson’s disease [26]. Moreover, an elevated soluble CD14 precursor has previously been found in CSF of patients with both Alzheimer’s disease and Parkinson’s disease in comparison to healthy controls [28].

The potential of peripheral (plasma) CD14 to serve as a biomarker for neurological conditions was confirmed by a meta-analysis, which found that elevated plasma soluble CD14 is associated with an increased risk of incident dementia as well as with markers of brain injury and aging, such as brain atrophy and executive function decline [29]. Our group recently employed a proteomic approach to explore biomarkers of neurodegenerative disorders in peripheral blood mononuclear cells (PBMCs) and again identified CD14 as a potential biomarker (manuscript in preparation).

In CSF of patients with Parkinson’s disease, elevated osteopontin levels were detected by the proteomic screening assay and validated using a quantitative ligand-binding assay. However, in plasma of patients with Parkinson’s disease, a significant increase of osteopontin concentration was only observed with the quantitative immunoassay, but not with proteomic screening. One reason for that finding may be the fact that osteopontin levels were 10-fold lower in plasma than in CSF samples. Thus, the screening assay may not have been sensitive enough to accurately measure osteopontin levels in plasma.

Osteopontin is a multifunctional glycophosphoprotein, which is involved in cell viability, oxidative stress/apoptosis, wound healing, inflammation, neurodegeneration, and tumor progression [30,31]. It has been implicated in promoting both tissue damage and repair mechanisms, and this pleiotropy may be partially related to the existence of different functional domains [32]. Osteopontin is expressed throughout various tissues, including the central nervous system [31].

As osteopontin has been implicated in biological processes involved in the pathogenesis of Parkinson’s disease, such as oxidative stress, mitochondrial dysfunction, cytokine regulation, and apoptosis, it has been hypothesized to play a role in the development of Parkinson’s disorder [33]. Previous studies have shown increased osteopontin levels in biological fluids from patients with Parkinson’s disease, even though there are certain inconsistencies among their findings. Maetzler et al. demonstrated elevated osteopontin concentrations in serum and CSF samples from patients with Parkinson’s disease [33]. However, Lin et al. found increased plasma osteopontin levels in patients with Parkinson’s disease, but no changes in their CSF osteopontin levels. Further, in the study by Lin et al., plasma osteopontin concentrations were correlated with C-reactive protein levels, motor impairment, and disease stage in patients with Parkinson’s disease [34].

In addition, in the current study, the quantitative osteopontin ligand-binding assay found that osteopontin levels were significantly increased in plasma and significantly decreased in CSF of patients with multiple sclerosis compared to healthy controls. Again, insufficient sensitivity of the proteomic osteopontin assay might have been a reason why such changes could not be identified. Notably, a previous meta-analysis summarizing 22 publications found elevated concentrations of osteopontin in both peripheral blood and CSF samples from patients with multiple sclerosis in comparison to controls [35]. This discrepancy cannot be definitively explained but may be related to the small sample size in our study. As osteopontin is also expressed in activated T-cells, higher osteopontin concentrations in body fluids suggest active inflammation, which is the main causative factor of multiple sclerosis [35].

In this study, the bioanalytes CD14 and osteopontin were identified by the scioDiscover proteomic screening assay and validated by quantitative fluorescence-based immunoassays. Based on the findings of the proteomic screening, vascular endothelial growth factor 165b (VEGF-165b), cancer antigen 15-3 (CA15-3), and cell adhesion molecule 1, 3, 5, 6, 8 (CEACAM-1, 3, 5, 6, 8) were also considered to be promising candidates; however, no quantitative ligand-binding assay with the required sensitivity was available for their validation.

The current study has several limitations. The sample size was relatively small, indicating limited power of the study; thus, a large-scale analysis would be required in the future to increase the reliability of the data and statistical analysis. In addition, only limited demographic and clinical information was available about the analyzed samples. Future studies should be conducted with well-characterized samples with known age, sex, and comorbidities. Finally, amyotrophic lateral sclerosis samples were analyzed only with quantitative ligand-binding assays but were not screened with the proteomic array.

## Conclusions

In conclusion, the current study demonstrated that a proteomic antibody microarray and a quantitative ligand-binding assay can be successfully combined to identify biomarker candidates in patients with neurodegenerative disorders. Elevated CD14 levels in CSF from patients with Alzheimer’s disease and elevated osteopontin levels in CSF from patients with Parkinson’s disease were detected using both the screening and confirmatory assays, rendering them promising biomarker candidates. The up- or downregulation of these proteins in the presence of the respective diseases could be linked to their pathogenesis. Larger-scale studies using this combined proteomic antibody microarray and quantitative ligand-binding assay approach should be employed to expand these findings.

## Supplementary Materials

Supplementary Table S1. Characteristics of the CSF samples from the validation cohort; Supplementary Table S2. Characteristics of the plasma samples from the validation cohort; Supplementary Table S3. Most differentially expressed proteins (p < 1e-02) identified by an antibody microarray in CSF of patients with neurodegenerative diseases in comparison to controls; Supplementary Table S4. Most differentially expressed proteins (p < 1e-02) identified by an antibody microarray in plasma of patients with neurodegenerative diseases in comparison to controls.

## Author Contributions

C.B., M.R., and M.S. designed the research; A.K. and R.S. performed the research; A.K., R.S., C.B., and M.S. analyzed the data; C.B., M.R., and M.S. wrote and edited the paper.

## Institutional Review Board Statement

The study was conducted in accordance with the Declaration of Helsinki, and samples were provided by commercial sources, which had obtained them under IRB-approved protocols (approved by WCG IRB Review Solutions with protocol code 20172667 and the Freiburg Ethics Commission International with protocol code 014/1616).

## Informed Consent Statement

Informed consent was obtained from all subjects involved in the study.

## Data Availability Statement

All other data used in the manuscript can be made available by the authors upon request.

## Acknowledgments

We thank Dr. Zoya Marinova for manuscript preparation.

## References

1. Erkkinen, M.G.; Kim, M.O.; Geschwind, M.D. Clinical neurology and epidemiology of the major neurodegenerative diseases. Cold Spring Harb Perspect Biol. 2018, 10, a033118.

2. Deuschl, G.; Beghi, E.; Fazekas, F.; Varga, T.; Christoforidi, K.A.; Sipido, E.; Bassetti, C.L.; Vos, T.; Feigin, V.L. The burden of neurological diseases in Europe: an analysis for the Global Burden of Disease Study 2017. Lancet Public Health. 2020, 5, e551–e567.

3. Tondo, G.; De Marchi, F. From biomarkers to precision medicine in neurodegenerative diseases: Where are we?. J Clin Med. 2022, 11, 4515.

4. Dugger, B.N.; Dickson, D.W. Pathology of neurodegenerative diseases. Cold Spring Harb Perspect Biol. 2017, 9, a028035.

5. Chaudhuri, A. Multiple sclerosis is primarily a neurodegenerative disease. J Neural Transm (Vienna). 2013, 120, 1463–1466.

6. Bourdenx, M.; Koulakiotis, N.S.; Sanoudou, D.; Bezard, E.; Dehay, B.; Tsarbopoulos, A. Protein aggregation and neurodegeneration in prototypical neurodegenerative diseases: Examples of amyloidopathies, tauopathies and synucleinopathies. Prog Neurobiol. 2017, 155, 171–193.

7. Kovacs, G.G. Molecular pathological classification of neurodegenerative diseases: turning towards precision medicine. Int J Mol Sci. 2016, 17, 189.

8. Golde, T.E. The therapeutic importance of understanding mechanisms of neuronal cell death in neurodegenerative disease. Mol Neurodegener. 2009, 4, 8.

9. Parnetti, L.; Gaetani, L.; Eusebi, P.; Paciotti, S.; Hansson, O.; El-Agnaf, O.; Mollenhauer, B.; Blennow, K.; Calabresi, P. CSF and blood biomarkers for Parkinson’s disease. Lancet Neurol. 2019, 18, 573–586.

10. Hansson, O. Biomarkers for neurodegenerative diseases. Nat Med. 2021, 27, 954–963.

11. Trejo-Lopez, J.A.; Yachnis, A.T.; Prokop, S. Neuropathology of Alzheimer’s disease. Neuro-therapeutics. 2022, 19, 173–185.

12. Koníčková, D.; Menšíková, K.; Tučková, L.; Hényková, E.; Strnad, M.; Friedecký, D.; Stejskal, D.; Matěj, R.; Kaňovský, P. Biomarkers of neurodegenerative diseases: biology, taxonomy, clinical relevance, and current research status. Biomedicines. 2022, 10, 1760.

13. Alcolea, D.; Beeri, M.S.; Rojas, J.C.; Gardner, R.C.; Lleó, A. Blood biomarkers in neurodegenerative diseases: implications for the clinical neurologist. Neurology. 2023, 101, 172–180.

14. Kwon, E.H.; Tennagels, S.; Gold, R.; Gerwert, K.; Beyer, L.; Tönges, L. Update on CSF biomarkers in Parkinson’s disease. Biomolecules. 2022, 12, 329.

15. Sill, M.; Schröder, C.; Shen, Y.; Marzoq, A.; Komel, R.; Hoheisel, J.D.; Nienhüser, H.; Schmidt, T.; Kastelic, D. Protein profiling gastric cancer and neighboring control tissues using high-content antibody microarrays. Microarrays (Basel). 2016, 5, 19.

16. Borrebaeck, C.A.; Wingren, C. Antibody array generation and use. Methods Mol Biol. 2014, 1131, 563–571.

17. Pušić, M.; Klančič, T.; Knific, T.; Vogler, A.; Schmidt, R.; Schröder, C.; Lanišnik Rižner, T. Antibody arrays identified cycle-dependent plasma biomarker candidates of peritoneal endometriosis. J Pers Med. 2022, 12, 852.

18. Janša, V.; Klančič, T.; Pušić, M.; Klein, M.; Vrtačnik Bokal, E.; Ban Frangež, H.; Rižner, T.L. Proteomic analysis of peritoneal fluid identified COMP and TGFBI as new candidate biomarkers for endometriosis. Sci Rep. 2021, 11, 20870.

19. Hufnagel, K.; Fathi, A.; Stroh, N.; Klein, M.; Skwirblies, F.; Girgis, R.; Dahlke, C.; Hoheisel, J.D.; Lowy, C.; Schmidt, R.; Griesbeck, A.; Merle, U.; Addo, M.M.; Schröder, C. Discovery and systematic assessment of early biomarkers that predict progression to severe COVID-19 disease. Commun Med (Lond). 2023, 3, 51.

20. Aldo, P.; Marusov, G.; Svancara, D.; David, J.; Mor, G. Simple Plex(™): A novel multi-analyte, automated microfluidic immunoassay platform for the detection of human and mouse cytokines and chemokines. Am J Reprod Immunol. 2016, 75, 678–693.

21. Kelley, S.L.; Lukk, T.; Nair, S.K.; Tapping, R.I. The crystal structure of human soluble CD14 reveals a bent solenoid with a hydrophobic amino-terminal pocket. J Immunol. 2013, 190, 1304–1311.

22. Wu, Z.; Zhang, Z.; Lei, Z.; Lei, P. CD14: Biology and role in the pathogenesis of disease. Cytokine Growth Factor Rev. 2019, 48, 24–31.

23. Reed-Geaghan, E.G.; Savage, J.C.; Hise, A.G.; Landreth, G.E. CD14 and toll-like receptors 2 and 4 are required for fibrillar A{beta}-stimulated microglial activation. J Neurosci. 2009, 29, 11982–11992.

24. Fassbender, K.; Walter, S.; Kühl, S.; Landmann, R.; Ishii, K.; Bertsch, T.; Stalder, A.K.; Muehlhauser, F.; Liu, Y.; Ulmer, A.J.; Rivest, S.; Lentschat, A.; Gulbins, E.; Jucker, M.; Staufenbiel, M.; Brechtel, K.; Walter, J.; Multhaup, G.; Penke, B.; Adachi, Y.; Hartmann, T.; Beyreuther, K. The LPS receptor (CD14) links innate immunity with Alzheimer’s disease. FASEB J. 2004, 18, 203–205.

25. Liu, Y.; Walter, S.; Stagi, M.; Cherny, D.; Letiembre, M.; Schulz-Schaeffer, W.; Heine, H.; Penke, B.; Neumann, H.; Fassbender, K. LPS receptor (CD14): a receptor for phagocytosis of Alzheimer’s amyloid peptide. Brain. 2005, 128, 1778–1789.

26. Letiembre, M.; Liu, Y.; Walter, S.; Hao, W.; Pfander, T.; Wrede, A.; Schulz-Schaeffer, W.; Fassbender, K. Screening of innate immune receptors in neurodegenerative diseases: a similar pattern. Neurobiol Aging. 2009, 30, 759–768.

27. Busse, S.; Hoffmann, J.; Michler, E.; Hartig, R.; Frodl, T.; Busse, M. Dementia-associated changes of immune cell composition within the cerebrospinal fluid. Brain Behav Immun Health. 2021, 14, 100218.

28. Yin, G.N.; Jeon, H.; Lee, S.; Lee, H.W.; Cho, J.Y.; Suk, K. Role of soluble CD14 in cerebrospinal fluid as a regulator of glial functions. J Neurosci Res. 2009, 87, 2578–2590.

29. Pase, M.P.; Himali, J.J.; Beiser, A.S.; DeCarli, C.; McGrath, E.R.; Satizabal, C.L.; Aparicio, H.J.; Adams, H.H.H.; Reiner, A.P.; Longstreth, W.T. Jr.; Fornage, M.; Tracy, R.P.; Lopez, O.; Psaty, B.M.; Levy, D.; Seshadri, S.; Bis, J.C. Association of CD14 with incident dementia and markers of brain aging and injury. Neurology. 2020, 94, e254–e266.

30. Icer, M.A.; Gezmen-Karadag, M. The multiple functions and mechanisms of osteopontin. Clin Biochem. 2018, 59, 17–24.

31. Rosmus, D.D.; Lange, C.; Ludwig, F.; Ajami, B.; Wieghofer, P. The role of osteopontin in microglia biology: current concepts and future perspectives. Biomedicines. 2022, 10, 840.

32. Cappellano, G.; Vecchio, D.; Magistrelli, L.; Clemente, N.; Raineri, D.; Barbero Mazzucca, C.; Virgilio, E.; Dianzani, U.; Chiocchetti, A.; Comi, C. The Yin-Yang of osteopontin in nervous system diseases: damage versus repair. Neural Regen Res. 2021, 16, 1131–1137.

33. Maetzler, W.; Berg, D.; Schalamberidze, N.; Melms, A.; Schott, K.; Mueller, J.C.; Liaw, L.; Gasser, T.; Nitsch, C. Osteopontin is elevated in Parkinson’s disease and its absence leads to reduced neurodegeneration in the MPTP model. Neurobiol Dis. 2007, 25, 473–482.

34. Lin, Y.; Zhou, M.; Dai, W.; Guo, W.; Qiu, J.; Zhang, Z.; Mo, M.; Ding, L.; Ye, P.; Wu, Y.; Zhu, X.; Wu, Z.; Xu, P.; Chen, X. Bone-derived factors as potential biomarkers for Parkinson’s disease. Front Aging Neurosci. 2021, 13, 634213.

35. Agah, E.; Zardoui, A.; Saghazadeh, A.; Ahmadi, M.; Tafakhori, A.; Rezaei N. Osteopontin (OPN) as a CSF and blood biomarker for multiple sclerosis: A systematic review and meta-analysis. PLoS One. 2018, 13, e0190252.

