## Supplementary Materials for "Exploration of novel biomarkers for neurodegenerative diseases using proteomic analysis and ligand-binding assays"

**Supplementary Table S1.** Characteristics of the CSF samples from the validation cohort.

| CSF | Control | AD | PD | MS | ALS |
|---|---|---|---|---|---|
| N | 84 | 18 | 35 | 10 | 14 |
| Age (mean±SD, years) | 56.9±17.5 | 68.8±8.05 | 70.6±7.72 | 46.0±5.66 | 68.0±8.86 |
| Sex (F/M/n.a.) | 48/26/10 | 6/4/8 | 14/16/5 | 1/1/8 | 2/12/0 |

AD: Alzheimer's disease; ALS: amyotrophic lateral sclerosis; CSF: cerebrospinal fluid; F: female; M: male; MS: multiple sclerosis; n.a.: not applicable; PD: Parkinson's disease; SD: standard deviation.

**Supplementary Table S2.** Characteristics of the plasma samples from the validation cohort.

| Plasma | Control | AD | PD | MS | ALS |
|---|---|---|---|---|---|
| N | 11 | 14 | 13 | 9 | 5 |
| Age (mean±SD, years) | n.a. | 85.5±4.37 | 77.5±8.43 | 57 | 52.8±10.64 |
| Sex (F/M/n.a.) | 0/0/11 | 6/0/8 | 4/4/5 | 1/0/8 | 0/5/0 |

AD: Alzheimer's disease; ALS: amyotrophic lateral sclerosis; F: female; M: male; MS: multiple sclerosis; n.a.: not applicable; PD: Parkinson's disease; SD: standard deviation.

**Supplementary Table S3.** Most differentially expressed proteins (p < 1e-02) identified by an antibody microarray in CSF of patients with neurodegenerative diseases in comparison to controls.

| Neurodegenerative disease | Protein | Antibody | logFC | Adj. P Val |
|---|---|---|---|---|
| **Alzheimer's disease** | LYVE1 | ab2405 | 0.97 | 1.6e-06 |
| | VCAM1 | ab1792 | 0.79 | 4.6e-05 |
| | CYTB | ab1241 | 0.75 | 4.6e-05 |
| | TIMP1 | ab1057 | 0.73 | 4.6e-05 |
| | MERTK | ab2456 | 1.56 | 6.5e-05 |
| | VEGF165b | ab2464 | -1.76 | 1.2e-04 |
| | MUC1 | ab1087 | -3.24 | 1.9e-04 |
| | GRN | ab2704 | 1.32 | 2.1e-04 |
| | MTOR | ab1153 | -1.41 | 2.8e-04 |
| | IL1AP | ab1770 | 0.88 | 4.8e-04 |
| | MUC17 | ab1267 | -1.42 | 6.6e-04 |
| | TNR1B | ab2455 | 0.67 | 7.7e-04 |
| | TIMP1 | ab1842 | 0.80 | 9.9e-04 |
| | UROM | ab2694 | -0.91 | 9.9e-04 |
| | SLAF8 | ab2129 | -1.13 | 9.9e-04 |
| | IBP4 | ab0483 | 0.61 | 1.2e-03 |
| | 2A5D | ab1225 | -1.65 | 1.2e-03 |
| | RET4 | ab1950 | 0.90 | 2.5e-03 |
| | CFAD | ab1219 | 0.76 | 2.5e-03 |
| | TNR14 | ab1765 | 0.57 | 2.5e-03 |
| | HMGB1 | ab1215 | -1.24 | 2.5e-03 |
| | CO2 | ab1216 | 0.78 | 2.6e-03 |
| | MPIP3 | ab1180 | -1.65 | 2.6e-03 |
| | HGFA | ab2253 | 0.72 | 2.7e-03 |
| | IBP6 | ab1979 | 0.58 | 2.9e-03 |
| | ANFB | ab0774 | -1.04 | 3.0e-03 |
| | PLK1 | ab1072 | -1.25 | 3.0e-03 |
| | CAMP | ab1184 | 1.15 | 3.4e-03 |
| | COMP | ab2307 | 0.76 | 3.5e-03 |
| | TGFR2 | ab2475 | 0.74 | 3.5e-03 |

| | | | | |
|---|---|---|---|---|
| | IFG2 | ab0482 | 0.59 | 3.5e-03 |
| | CXCR5 | ab1059 | -1.13 | 3.5e-03 |
| | GPI8 | ab1317 | -1.31 | 3.5e-03 |
| | ACY1 | ab2821 | 0.62 | 3.9e-03 |
| | PARK7 | ab2223 | 0.75 | 4.3e-03 |
| | TNPO3 | ab0583 | -1.11 | 4.3e-03 |
| | CATB | ab1172 | -1.30 | 4.3e-03 |
| | CD28 | ab1559 | -1.29 | 5.4e-03 |
| | RARR2 | ab2130 | -0.93 | 5.8e-03 |
| | CD14 | ab1393 | 0.95 | 5.9e-03 |
| | CTNB1 | ab1183 | -1.52 | 6.2e-03 |
| | ING1 | ab1032 | -0.95 | 6.8e-03 |
| | BEX3 | ab0714 | -0.93 | 7.0e-03 |
| | AOXA | ab0441 | -1.27 | 7.0e-03 |
| | TMM54 | ab0075 | 2.39 | 7.1e-03 |
| | SODC | ab0644 | -0.90 | 7.3e-03 |
| | CAD13 | ab1230 | -1.09 | 7.3e-03 |
| | IL26 | ab1328 | -1.28 | 7.3e-03 |
| | G3P | ab1004 | -1.37 | 7.3e-03 |
| | RENI | ab2133 | -1.67 | 7.3e-03 |
| | A1BG | ab2811 | 0.72 | 7.4e-03 |
| | CD38 | ab1537 | 1.80 | 8.1e-03 |
| | ANGI | ab1743 | 0.92 | 8.1e-03 |
| | IGF1R | ab1995 | -1.21 | 8.1e-03 |
| | UB2R1 | ab1305 | -1.27 | 8.5e-03 |
| | S10A8/9 | ab1624 | 1.43 | 9.2e-03 |
| | HAVR2 | ab2067 | -1.37 | 9.2e-03 |
| | G3P | ab1332 | -1.36 | 9.8e-03 |
| **Parkinson's disease** | OSTP | ab1737 | 0.83 | 3.5e-03 |

**Supplementary Table S4.** Most differentially expressed proteins (p < 1e-02) identified by an antibody microarray in plasma of patients with neurodegenerative diseases in comparison to controls.

| Neurodegenerative disease | Protein | Antibody | logFC | Adj. P Val |
|---|---|---|---|---|
| **Alzheimer's disease** | PGH2 | ab0492 | 2.95 | 1.3e-03 |
| | CORIN | ab1954 | 1.84 | 6.1e-03 |
| **Parkinson's disease** | CD14 | ab1393 | 1.43 | 3.4e-04 |
| | TOP1 | ab1298 | 1.69 | 8.6e-04 |
| | IgE | ab1506 | 1.29 | 8.6e-04 |
| | BASI | ab1487 | 1.30 | 2.4e-03 |
| | Fc fusion with TNR1B | ab0520 | 1.63 | 2.6e-03 |
| | ONCM | ab1885 | 0.97 | 3.6e-03 |
| | CD8A | ab1376 | 1.31 | 4.7e-03 |
| | CEAM6 | ab1633 | 1.25 | 4.7e-03 |
| | JAK2 | ab0978 | 1.19 | 4.7e-03 |
| | TNR1A | ab2332 | 1.11 | 4.7e-03 |
| | IL13 | ab1716 | -1.25 | 4.7e-03 |
| | CCCL7 | ab1695 | -1.00 | 5.2e-03 |
| | MUC1 | ab1087 | 1.73 | 5.6e-03 |

| | | | | |
|---|---|---|---|---|
| | IBP2 | ab1835 | 1.09 | 5.6e-03 |
| | S10A8/9 | ab2724 | 0.88 | 5.6e-03 |
| | TR13B | ab2250 | 1.20 | 6.1e-03 |
| | IgE | ab1593 | 1.03 | 8.4e-03 |
| | CD44 | ab1437 | 1.00 | 8.3e-03 |
| **Multiple sclerosis** | ANGT | ab2292 | 1.79 | 1.9e-04 |
| | MMP9 | ab1946 | 1.15 | 8.0e-04 |
| | TR13B | ab2250 | 1.31 | 5.5e-03 |
| | VEGF165b | ab2464 | 1.18 | 1.2e-03 |
